# Eccentric exercise ≠ eccentric contraction

**DOI:** 10.1101/2023.11.23.568422

**Authors:** Paolo Tecchio, Brent J. Raiteri, Daniel Hahn

## Abstract

Whether eccentric exercise involves active fascicle stretch is unclear due to muscle-tendon unit (MTU) series elasticity. Therefore, this study investigated the impact of changing the activation timing and level (i.e., pre-activation) on muscle fascicle kinematics and kinetics of the human tibialis anterior during dynamometer-controlled maximal voluntary MTU-stretch-hold contractions. B-mode ultrasound and surface electromyography were employed to assess muscle fascicle kinematics and muscle activity levels, respectively. While joint kinematics were similar for MTU-stretch-hold contractions, increasing pre-activation increased fascicle shortening and stretch amplitudes (9.9 - 23.2 mm, *p* ≤ 0.015). This led to increasing positive and negative fascicle work with increasing pre-activation. Despite significantly different fascicle kinematics, similar peak fascicle forces during stretch occurred at similar fascicle lengths and joint angles regardless of pre-activation. Similarly, residual force enhancement (rFE) following MTU stretch was not significantly affected (6.5–7.6 %, *p* = 0.559) by pre-activation, but rFE was strongly correlated with peak fascicle force during stretch (*r*_rm_ = 0.62, *p* = 0.003). These findings highlight that apparent eccentric exercise causes shortening-stretch contractions at the fascicle level rather than isolated eccentric contractions. The constant rFE despite different fascicle kinematics and kinetics suggests that a passive element was engaged at a common muscle length among conditions (e.g., optimal fascicle length). Although it remains unclear whether different fascicle mechanics trigger different adaptations to eccentric exercise, this study emphasizes the need to consider MTU series elasticity to better understand the mechanical drivers of adaptation to exercise.

**New & Noteworthy:** Apparent eccentric exercises do not result in isolated eccentric contractions, but shortening-stretch contractions at the fascicle level. The amount of fascicle shortening and stretch depend on the pre-activation during the exercise and *cannot* be estimated from the muscle-tendon unit or joint kinematics. As different fascicle mechanics might trigger different adaptations to eccentric exercise, muscle-tendon unit series elasticity and muscle pre-activation need to be considered when eccentric exercise protocols and designed and evaluated.

## Introduction

Muscles actively change length to power everyday movement. But despite being the motors of movement, muscles function surprisingly better as brakes than motors (1). Accordingly, muscle forces are enhanced during and following active muscle stretch (i.e., eccentric contraction) compared with contractions at a constant length (i.e., isometric contraction) or during active muscle shortening (i.e., concentric contraction) when muscle activity is matched (2–5). Accordingly, producing a given force eccentrically requires less muscle activity compared with an isometric or a concentric contraction. Since active muscle volume theoretically determines energy expenditure (6), the neuromuscular efficiency of eccentric contractions is therefore increased (7, 8).

As eccentric contractions require less energy and muscle activity for equivalent force production, eccentric exercise has received considerable attention over the last decades to rehabilitate injuries, to improve athletic performance, and to help manage neuromuscular diseases (9–13). Accordingly, many training studies aimed to determine the specific adaptations of the muscle-tendon unit (MTU) to eccentric exercise in comparison to concentric exercise. The specific adaptations to eccentric exercise primarily include an increase in fascicle length (14). However, the fascicle length increases following eccentric exercise are highly variable among studies. By examining 15 in vivo human studies on training adaptations to eccentric exercise (11, 15–28), we found that fascicle length changed from −3% (i.e., decreased) to 34% (i.e., increased, see Figure S1). Accordingly, it is important to question why the fascicle length changes to eccentric exercise are so variable.

One factor that could affect the muscle lengths and forces attained during and following eccentric contractions and the subsequent MTU adaptations following eccentric exercise is the pre-activation timing and level before MTU stretch. Because of the typical arrangement of a distal elastic tendon in series with a muscle within a lower limb MTU, the length changes of muscle and tendon can be decoupled (29, 30). For example, a muscle can still shorten against the elastic tendon while the MTU is stretched when there is no pre-activation. Accordingly, apparent eccentric exercises (i.e., MTU lengthening exercises) do not necessarily result in isolated eccentric contractions, but can result in a combination of muscle contraction types. Therefore, the highly variable fascicle length changes to apparent eccentric exercise could be due to different muscle fascicle behaviors during these exercises.

During apparent eccentric exercise, the decoupling of muscle and tendon length changes can also affect the fascicle stretch amplitudes (31, 32). Differences in fascicle stretch amplitude during eccentric exercises might affect peak fascicle force during stretch, the residual force enhancement (rFE) following stretch (33–35), and the active and passive contributions to eccentric force production. This is because rFE was found to be related to the peak force and/or torque during stretch (33, 36, 37). However, it is unclear whether peak force is affected more by stretch amplitude or fascicle length (38). It is also unclear whether the rFE following active muscle stretch is affected more by the stretch amplitude or the final muscle length. Animal studies showed that rFE increased with increasing muscle stretch amplitude up until a point when stretching occurred at lengths where passive muscle force naturally existed (2, 39). A more recent in vivo study on the human knee extensors however showed that rFE primarily depends on the final fascicle length rather than the fascicle stretch amplitude during submaximal MTU-stretch-hold contractions (40). These findings are potentially relevant for the specific adaptations to eccentric exercise as rFE is thought to be related to the properties of the giant muscle protein titin. In turn, changes in titin stiffness have been reported to drive longitudinal muscle hypertrophy (41), which could be another potential reason for the highly variable fascicle length adaptations to eccentric exercise reported in the literature.

Therefore, the aim of this research was to study the effect of MTU series elasticity on the fascicle kinematics (i.e., behavior) and kinetics (i.e., force and work) during apparent eccentric exercise. Changes in human tibialis anterior (TA) MTU series elasticity were induced by implementing three different pre-activation timings and levels (PLs) that were torque controlled (0, 50%, and 95% of the joint-angle-specific peak active torque) prior to MTU stretch. We hypothesized that both fascicle shortening and stretch amplitudes would increase with increasing PL because of a reduction in MTU series elasticity. Based on these expected fascicle kinematics, we also hypothesized that both the positive and negative fascicle work produced during the contractions would increase with increasing PL. We further hypothesized that similar peak forces during MTU stretch would occur at similar fascicle lengths following different fascicle stretch amplitudes that would be in accordance with TA’s active force-length relation (42, 43). Our final hypothesis was that rFE would not be related to fascicle stretch amplitude, but to the peak fascicle force achieved during MTU stretch. This is because energy storage within passive elements (e.g., titin) should be better reflected by peak fascicle force than fascicle stretch amplitude.

## Methods

### Participants

A simulation-based a-priori power analysis using Superpower (44) estimated that a sample size of eleven participants would be necessary to detect our smallest effect size of interest of 0.78 (based on pilot data of *n* = 4) with 80% power at a two-tailed alpha level of 5%. Therefore, twelve healthy participants (five women; age: 25.6 ± 2.7 years; mass: 71.6 ± 14.1 kg) gave free written informed consent prior to participating in this study. All participants reported no recent (<24 months) history of lower limb injuries or neuromuscular disorders. The experimental procedures were approved by the local Ethics Committee of the Faculty of Sport Science at Ruhr University Bochum (EKS V10/2022). The study was conducted in accordance with the ethical principles outlined within the Declaration of Helsinki with the exception that our study was not pre-registered.

### Experimental setup

Participants performed ankle dorsiflexion contractions with their right dorsiflexors while sitting in a reclined position (~70° from the horizontal) on the seat of a dynamometer (IsoMed2000, D&R Ferstl GmbH, Hemau, Germany). Each participant’s foot was fixed to the footplate attachment of the dynamometer by straps and a custom 3D-printed inverse U-shaped frame over the metatarsal joints (the stereolitography models are available at https://github.com/PaulT95/Spike2_Collecting/tree/main/FussHalter_Model). The neutral position (i.e., 0°) of the ankle joint was defined as an angle of 90° between the footplate and the tibia, which was achieved when both were at an angle of 45±1° relative to the horizontal (measured by a digital inclinometer, accuracy ±0.1°; Beaspire, Amazon, Seattle). The right thigh was supported by a cushioned dynamometer attachment to limit accessory movements due to hip flexor activation (43, 45). The ankle joint axis of rotation was aligned with the dynamometer axis of rotation during a maximal voluntary fixed-end contraction at 15° plantarflexion as this angle was at the midway point of the range of motion (ROM) of the eccentric contractions. The foot position was adjusted until a laser, which was projected from a laser pointer located within the dynamometer’s axis of rotation, was aligned on the skin with the transmalleolar axis. The transmalleolar axis was therefore assumed to correspond to the ankle joint’s axis of rotation. Further, the dynamometer was inverted by 5° relative to the vertical to supinate the foot and ensure the dynamometer and transmalleolar axes of rotation were approximately parallel. During the experiment, participants were asked to fold their arms across their chest prior to each contraction to limit accessory movements. Visual live feedback of the torque and the desired torque trace (see Experimental protocol) were provided on a screen that was located directly in front of the participants.

### Experimental protocol

The experiment consisted of two sessions. In the first session, participants were familiarized with performing maximal voluntary contractions (MVCs) on the dynamometer. The second session served as the main experimental session where the actual measurements were made. Both sessions started with a standardized warm-up that included 8-12 submaximal (~50–80% of perceived maximal effort) ankle dorsiflexion contractions at both short (−5° dorsiflexion, DF) and long (+35° plantarflexion, PF) MTU lengths to precondition the MTU (46, 47).

Following preconditioning, two brief (~3 s) maximal voluntary fixed-end contractions were performed at DF. If the difference between the peak-to-peak torques of the two contractions was greater than 5%, additional contractions were performed until the difference between the two strongest contractions was less than 5%. The strongest contraction of the two was used to determine three PL levels for the MTU-stretch-hold contractions, namely 0% (PL0), 50% (PL50), and 95% (PL95) of the joint-angle-specific maximal active torque. One brief (~3 s) maximal voluntary fixed-end contraction was also performed at PF to determine the slopes of the ramps (see Contraction conditions). Maximal voluntary fixed-end contractions at PF and the MTU-stretch-hold contractions with the three different PLs were then performed in a randomized order.

The MTU-stretch-hold contractions were performed over a ROM from the short position at DF to the long position at PF (*ω* = 40°/s, acc = 1000°/s^2^, dec = 400°/s^2^). At least two contractions per condition were conducted and if the mean electromyography (EMG) root-mean-square (RMS) amplitude (0.25 s overlapping window, see Figure 1) during the steady state differed by more than 10% between two fixed-end contractions at PF, additional fixed-end reference contractions were performed. Additional MTU-stretch-hold contractions were performed if the EMG RMS amplitude during the steady state differed by more than 20% between the MTU-stretch-hold contraction and the averaged value from the valid fixed-end reference contractions. However, in order to minimize fatigue, no more than four contractions per condition were performed to ensure that participants did not exceed a predetermined limit of 18 sustained MVCs during the experiment. To further minimize fatigue, a rest period of 3 min and 30 s up to 5 min was interposed between the sustained MVCs. Standardized verbal encouragement was provided during each MVC by the investigator (48).

**Figure 1.**
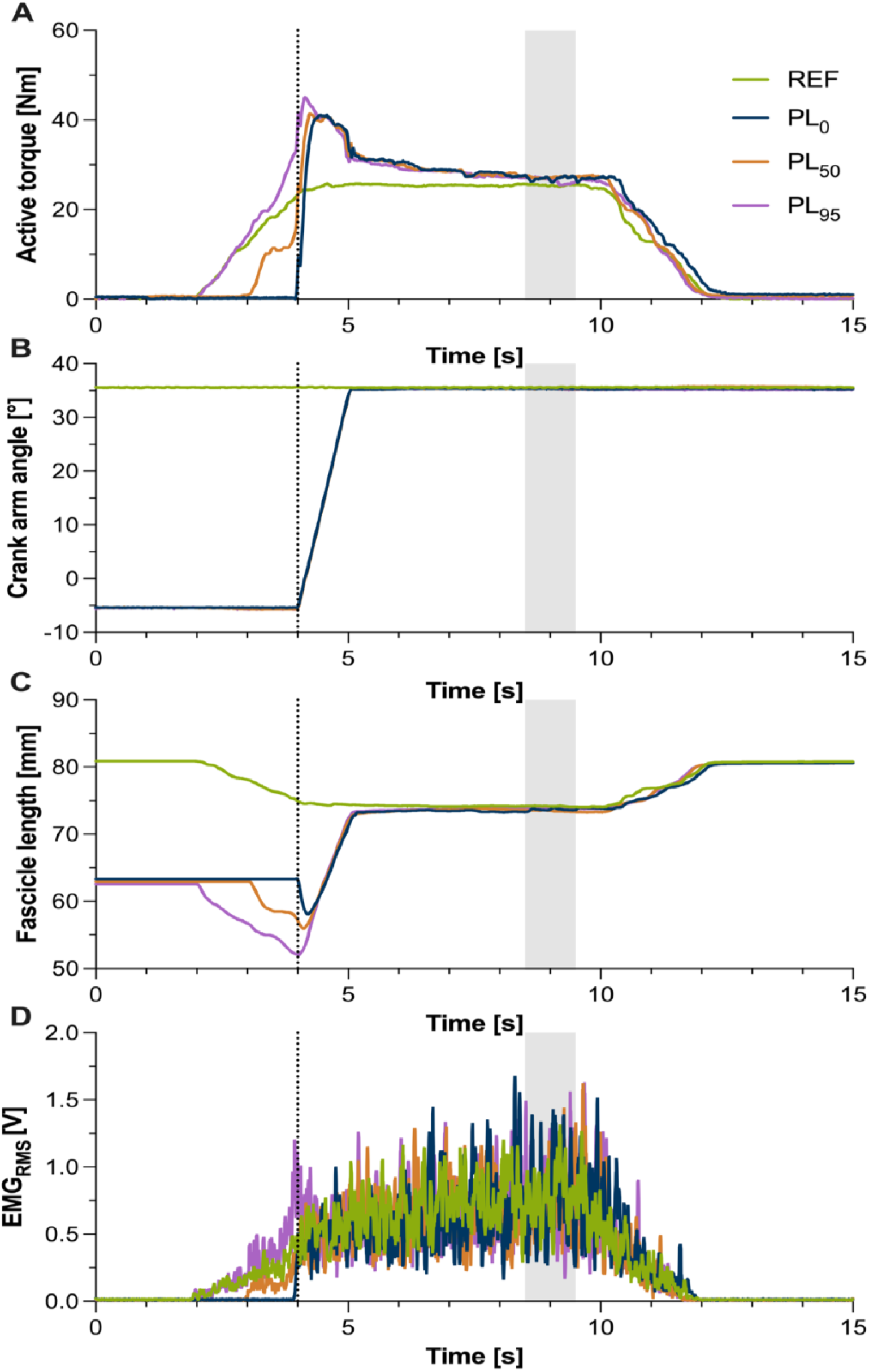
(**A**) Active torque-time traces, (**B**) crank-arm angle-time traces, (**C**) fascicle length-time traces, (**D**) and the centered root-mean-squared (RMS, 25 ms time window) amplitude-time traces of the surface electromyography (EMG) signal across the MTU-stretch-hold and fixed-end reference conditions from one participant. The green traces represent the fixed-end reference contraction at the long position (+35° plantarflexion). The blue, orange, and purple traces represent the MTU-stretch-hold conditions with a 0%, 50%, and 95% PL, respectively. The rotation of the footplate to induce stretch of the TA MTU was always automatically triggered at 4 s as indicated by the black vertical dotted line, and ended at 5 s (ω **=** 40 °/s). Positive crank arm angles indicate ankle plantar flexion. The steady state where residual force enhancement (rFE) was assessed is highlighted by the gray shaded area and spans from 3.5-4.5 s after the end of the rotation.

Following all MVCs, six passive ankle joint rotations were performed over the ROM from the DF to the PF positions and vice versa (*ω* = 10°/s, acc = 100°/s^2^, dec = 100°/s^2^). Passive ankle joint torques were also recorded statically at six different angles within the ROM to calculate a cubic polynomial fit for passive torque-angle data to allow net active ankle joint torques to be estimated post-testing.

### Contraction conditions

For the PL0 condition, participants were instructed to remain relaxed until the rotation of the dynamometer was triggered. In the PL50 and PL95 conditions, participants were instructed to gradually increased their voluntary dorsiflexion effort and corresponding dorsiflexion torque to reach the desired PL by following a superimposed torque ramp plotted on a screen in front of the them, which was required before the dynamometer rotation was triggered. The ramp rates (i.e., slopes) for PL50 and PL95 conditions were matched, with a duration of ~2 s to reach PL95 and 1 s to reach PL50. During all MTU-stretch-hold contractions, participants were instructed to fully activate their TA and to contract “as hard and fast as possible” as soon as the dynamometer rotation started. Two vertical dashed lines were plotted on the abovementioned screen at the onset of rotation and five seconds after the end of the rotation to indicate the duration participants were required to maximally contract before gradually relaxing.

### Surface electromyography (EMG)

Surface EMG (NeuroLog System NL905, Digitimer Ltd, UK) was used to record TA muscle activity. After skin preparation (shaving, abrading, cleaning the skin with antiseptic), two surface electrodes were attached over the distal muscle belly with an inter-electrode distance of ~20 mm. The electrodes were placed distally to the ultrasound transducer, which was located over the mid muscle belly. To ensure that the electrodes remained over the TA muscle tissue during the contractions, the muscle-tendon junction was visualized with ultrasound imaging during a maximal voluntary fixed-end contraction at DF, and a pen mark was drawn on the skin to indicate the most-distal location the electrodes could be placed.

### Ultrasound imaging (US)

TA muscle architecture was imaged using a PC-based ultrasound system (ArtUs EXT-1H, Telemed, Vilnius, Lithuania) and a flat, linear ultrasound transducer with 128 elements (LV8-5N60-A, B-mode, 8.0 MHz, 60 mm width and 50 mm depth, Telemed, Vilnius, Lithuania). The images were captured at 33 Hz (high line density) and the transducer was firmly attached over the mid muscle belly using a custom 3D-printed plastic frame and adhesive bandage.

### Data collection

Net ankle joint torque and dynamometer crank-arm angle were digitally sampled at 2 kHz and synchronized with the ultrasound system and the EMG signal using a 16-bit Power 1401 and Spike2 (v. 8.13) data collection system (Cambridge Electronic Design, UK). The ultrasound system was triggered to record frames automatically by square-wave pulses (80/20 duty cycle at 100 Hz, 4 V amplitude, 10 µs per clock tick). The dynamometer crank-arm rotation during the MTU-stretch-hold contractions was automatically triggered by a digital output from the 1401 device (the data acquisition script and the circuit schematic are available at https://github.com/PaulT95/Spike2_Collecting).

### Fascicle length tracking

Muscle fascicles of the TA’s superficial compartment were automatically tracked using an updated version of Ultratrack (49). Specifically, we implemented a Lukas-Kanade-Tomasi-based affine optic flow algorithm as in previous studies (40, 50). To limit subjectivity in the fascicle length determination, a representative fascicle was automatically detected and linearly extrapolated to find its intersections with the automatically-detected superficial and central aponeuroses of TA (examples are available in the GitHub repository). For quality control, the automated detection procedures were visually inspected and if necessary, manually corrected (*n* = 4). The fascicle angle was defined as the angle between the tracked fascicle and the horizontal. If the central aponeurosis angle was constantly greater than 5° relative to the horizontal throughout the contraction (*n* = 2), the fascicle angle values were manually corrected by manually tracking the central aponeurosis angle relative to the horizontal and then added to the fascicle angle.

### Data analysis

All data were post-processed using custom-written scripts written in MATLAB (R2021b). The data were cropped based on the times of the first and last digital synchronization pulses of the ultrasound recording. Fascicle length data was resampled at a frequency of 2 kHz. Torque, crank-arm angle, and fascicle length data were filtered using a zero-lag dual-pass-corrected (51) fourth-order 12 Hz low-pass Butterworth filter. The EMG signal was filtered using a dual-pass second-order 10-450 Hz band-pass Butterworth filter. The DC offset was then removed. Following this, the centered moving RMS amplitude was calculated over a 25 ms window. In addition, an envelope was obtained using a dual-pass corrected second-order low-pass 5 Hz Butterworth filter on the band-passed signal (52, pg. 189).

Reference (REF) EMG and active torque values were based on the two fixed-end contractions at PF with the smallest symmetrized percent difference (SPD) (53) in mean steady-state EMG RMS amplitude. The steady state was defined as the time period from 3.5 to 4.5 s after the end of the crank-arm rotation. If the differences in mean steady-state EMG RMS amplitude between all fixed-end contractions were within 10%, the participant’s data was included in the analysis. Subsequently, the MTU-stretch-hold contraction with the smallest mean steady-state EMG RMS amplitude percentage difference relative to the REF condition was included in the analysis; however, conditions were excluded when they did not fall within ±20% of the REF condition. Mean torque and mean fascicle length were also calculated during the steady state and rFE was calculated as the difference in mean steady-state active torque between the MTU-stretch-hold and REF conditions divided by the REF condition.

Active torque was determined as the difference between the recorded net ankle joint torque and the passive net ankle joint torque recorded when participants were instructed to relax (i.e., no muscle activity changes were visually detected). Joint work was calculated as the cumulative trapezoidal integral of active torque as a function of the crank-arm angle. TA tendon force was then estimated by dividing the active torque by the TA literature-based moment arm measured during MVC (53, see Figure S2) and this estimated net dorsiflexor force in the longitudinal direction was multiplied by 50% based on TA’s volume (60%) relative to the total dorsiflexor volume (55) and based on TA’s apparent 45-52% relative torque contribution during MVC (56). Fascicle force was then calculated by dividing TA tendon force by the cosine of the fascicle angle. Fascicle work was then estimated by multiplying fascicle force and fascicle displacement. Fascicle velocity was defined as the first centered derivative of fascicle length within a 4 ms time window.

Fascicle shortening amplitude was calculated by subtracting the passive fascicle length before the contraction onset and the shortest fascicle length detected as the minimum value during the contraction. The fascicle shortening amplitude also accounted for any passive fascicle stretch that occurred due to a slight delay in contraction onset relative to rotation onset in PL0. The fascicle stretch amplitude was determined by subtracting the mean fascicle length during the steady state and the shortest fascicle length. Further, the fascicle stretch amplitude to peak fascicle force was determined as the difference between the length where the peak fascicle force occurred during stretch and the shortest fascicle length.

The cumulative fascicle work in the shortening phase (positive work) and the stretch phase (negative work) were also calculated. Net fascicle work was then calculated as the sum of positive and negative work in the MTU-stretch-hold conditions. Further, peak fascicle force and the corresponding time during rotation were determined from the selected trials along with the active torque, the crank arm angle, the fascicle length, and the EMG envelope value at this time. Similarly, peak active torque during rotation was identified and the same variables as above were extracted at this time.

### Statistics

Statistical analysis was performed with Prism (GraphPad Prism 9.5.0, San Diego, California, USA) and RStudio (v4.22, 2022, Boston, Massachusetts, USA). A one-way repeated-measures analysis of variance (ANOVA) was used to identify significant mean differences in the variables of interest across the conditions (REF vs. MTU-stretch-hold conditions, or only among MTU-stretch-hold conditions). Dunnett and Tukey post hoc tests were used to analyze mean differences between conditions. Repeated-measures Pearson correlations were performed using the “*rmcorr package*” (57) to determine the common within-subject association between peak fascicle force and the fascicle stretch amplitude until peak fascicle force, as well as between rFE and the total fascicle stretch amplitude or peak fascicle force. The family-wise alpha level was set at 5% and Bonferroni-adjusted for the three repeated-measures correlations. Results are reported as mean ± standard deviation.

## Results

### EMG matching

Although the ANOVA revealed a significant main effect (−6.9 to −1.8%, *p* = 0.032) for the EMG RMS percentage difference relative to the REF, all values from the selected MTU-stretch-hold contractions fell within our pre-selected criteria of ±20% REF_EMG_.

### Fascicle and joint mechanics

The ankle joint and MTU kinematics (i.e., crank arm angle at DF, PF and angular velocity) showed no significant differences among conditions (*p* = 0.485, Figure S3). TA muscle fascicles initially shortened in the REF and MTU-stretch-hold contractions. Across the MTU-stretch-hold contractions, fascicle kinematics were clearly different during the rotation despite similar joint kinematics (Figure 2). The amount of fascicle shortening and stretch significantly increased (*p* ≤ 0.015) almost linearly as the PL increased among the MTU-stretch-hold contractions (9.9 ± 2.8 mm in PL0, 14.0 ± 4.8 mm in PL50, and 16.1 ± 3.2 mm in PL95, respectively for fascicle shortening; 16.0 ± 3.8 mm in PL0, 20.3 ± 4.7 mm in PL50, and 23.2 ± 4.3 mm in PL95, respectively for fascicle stretch; Figure 3A-B, Table 1).

**Figure 2.**
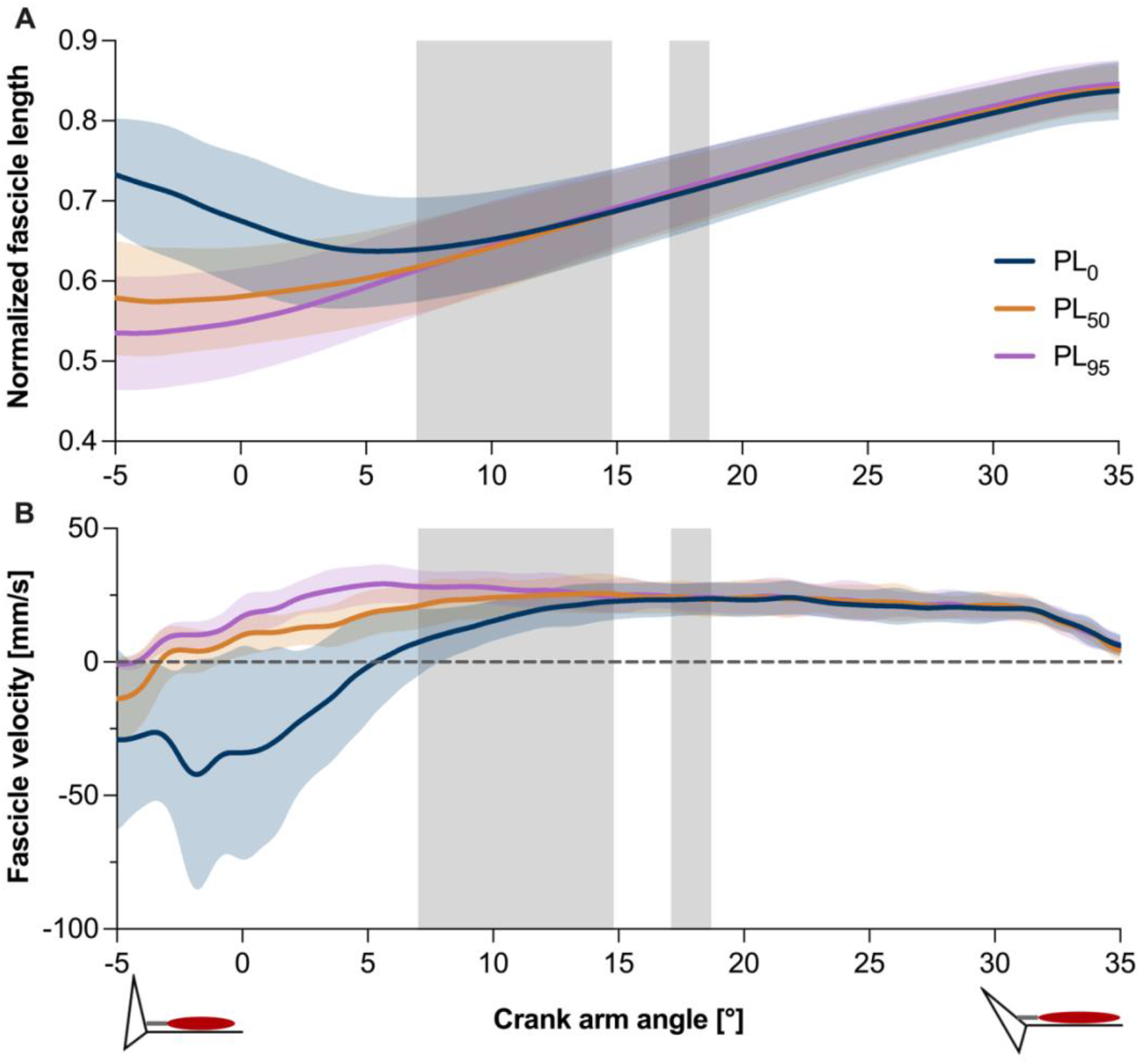
Normalized TA fascicle lengths (relative to the passive fascicle length at the long position, +35° plantarflexion) (**A**) and absolute TA fascicle velocities (**B**) across participants during the one-second crank-arm rotation for the three MTU-stretch-hold contractions (PL0, PL50, and PL95). The solid lines indicate the mean and the shaded areas indicate the standard deviation. The gray shaded areas indicate the range where the peak torque (on the left) and the peak fascicle force occurred (on the right) during the crank-arm rotation, respectively.

**Figure 3.**
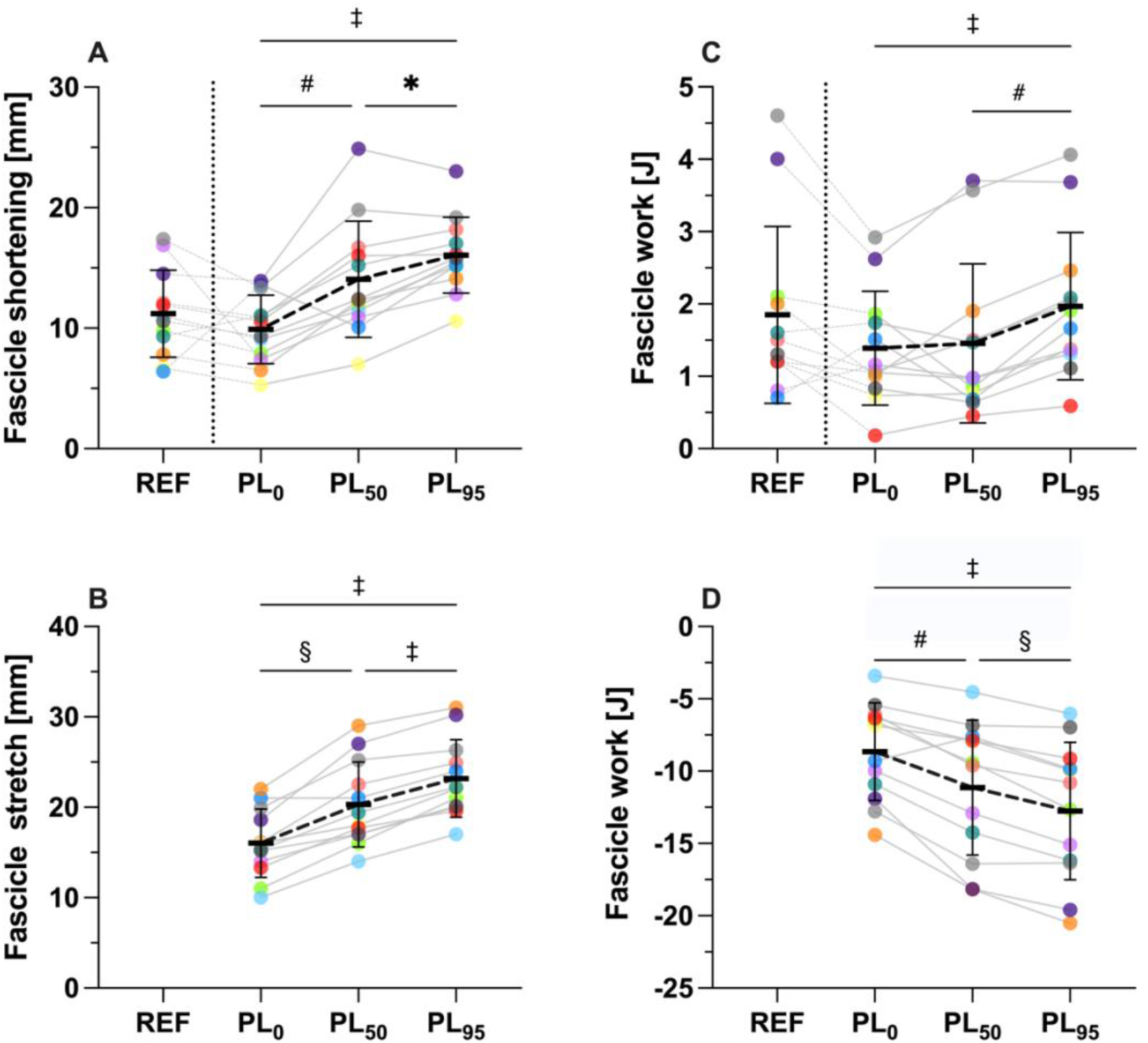
Individual (*n* = 12) and mean magnitudes of tibialis anterior fascicle shortening (A), fascicle stretch (B), positive fascicle work (C), and negative fascicle work (D) across the contraction conditions (REF, PL0, PL50 and PL95). Each colored dot represents a single participant. The thick and thin black horizontal and vertical bars indicate the group mean and standard deviation, respectively. * *p* <0.05; ^#^ *p* < 0.01; ^§^ *p* < 0.001; ^‡^ *p* < 0.0001.

**Table 1.** Fascicle kinematics, fascicle kinetics, and joint work among the different conditions.

|  | Fascicle kinematics and kinetics |  |  |  |  | Joint Work |
| --- | --- | --- | --- | --- | --- | --- |
|  | Fascicle length (mm) | Fascicle shortening (mm) | Fascicle stretch (mm) | Fascicle Positive Work (J) | Fascicle Negative Work (J) | Joint Work (J) |
| REF | 65.8 ± 11.4 | 11.2 ± 3.6 | - | 1.85 ± 1.22 | - | - |
| PL 0 | 65.2 ± 12.4 | 9.9 ± 2.8 | 16 ± 3.8 | 1.39 ± 0.79 | -8.66 ± 3.37 | 26.5 ± 7.2 |
| PL 50 | 65.4 ± 12 | 14 ± 4.8 | 20.3 ± 4.7 | 1.45 ± 1.1 | -11.14 ± 4.67 | 30.1 ± 8.7 |
| PL 95 | 65.7 ± 12.4 | 16.1 ± 3.2 | 23.2 ± 4.3 | 1.97 ± 1.02 | -12.76 ± 4.76 | 31.7 ± 8.5 |

Values are mean ± SD; N = 12. REF = reference, PL = pre-activation level.

Positive fascicle work was 1.9 ± 1.2 J in the REF condition. In the MTU-stretch-hold contractions, positive fascicle work was similar in PL0 (1.4 ± 0.8 J) and PL50 (1.5 ± 1.1 J, *p* = 0.0934), but significantly higher in PL95 (2.0 ± 1.0, *p* ≤ 0.003; Figure 3C, Table 1). Conversely, negative fascicle work linearly significantly increased (*p* ≤ 0.003) with increasing PL in the MTU-stretch-hold contractions (−8.7 ± 3.4 J in PL0, −11.1 ± 4.7 J in PL50, and −12.8 ± 4.8 J in PL95, respectively; Figure 3D and Table 1). In addition, active ankle joint work was significantly higher in PL50 (*p* = 0.003) and PL95 (*p* < 0.001) compared with PL0, but similar between PL50 and PL95 (*p* = 0.060) (for detailed results refer to Table 1).

### Peak fascicle force during stretch

TA peak fascicle force during rotation was similar among the MTU-stretch-hold contractions (579 ± 160 N; *p* = 0.645; Figure 5B) and it occurred at similar fascicle lengths (53.2 ± 11.4 mm; *p* = 0.757; Figure 5D) and velocities (23.6 ± 6.2 mm/s; *p* = 0.218; Figure 5D), independent of the PL. Muscle activity (i.e., EMG envelope), crank arm angle, and active torque were also similar among the MTU-stretch-hold contractions when peak fascicle force occurred (0.593 ± 0.214 V, 17.8 ± 4.6° and 47.6 ± 12.9 Nm, *p* ≥ 0.06 respectively; Figure 5).

**Figure 4.**
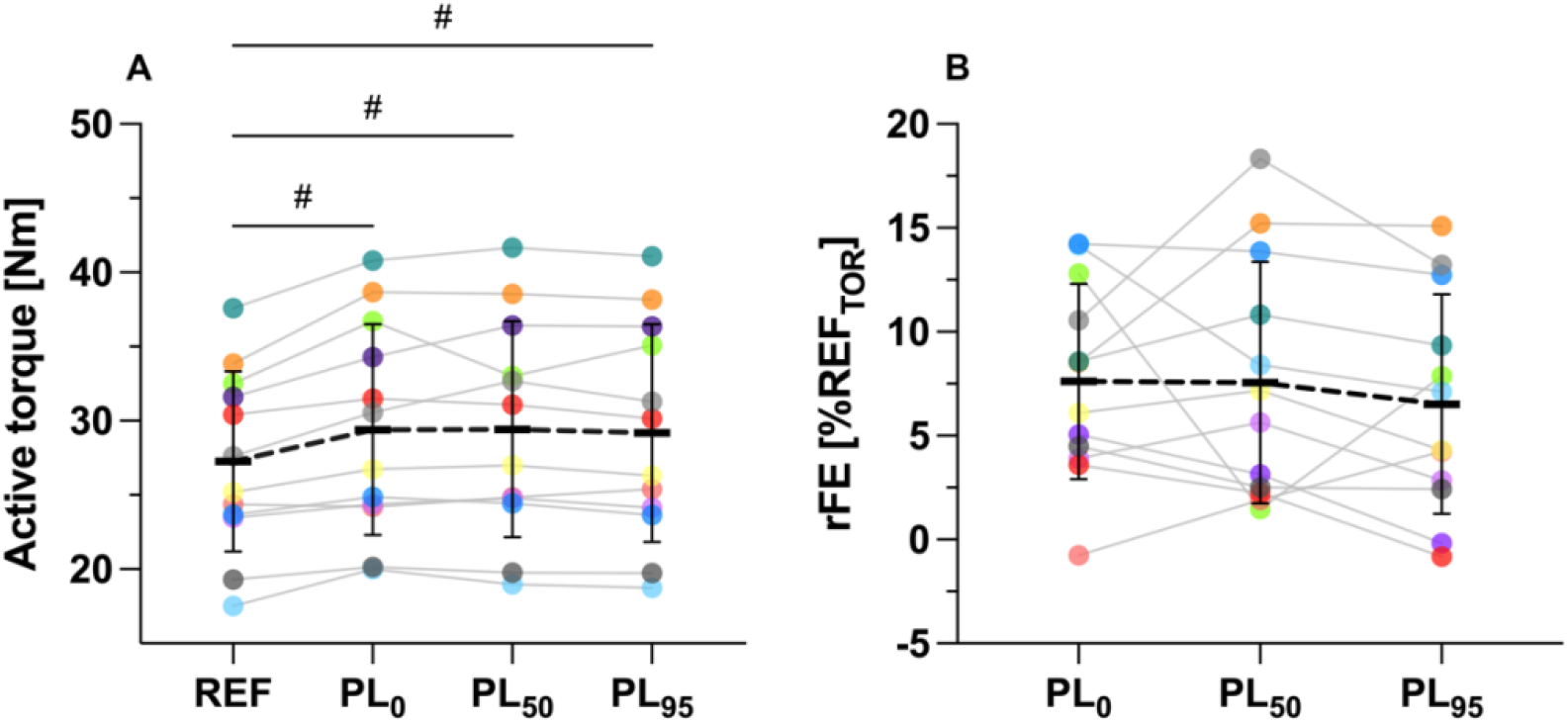
Individual (*n* = 12) and mean (A) net ankle joint torque values and (B) percentage residual force enhancement values at the long muscle length during the steady state phase of the fixed-end reference (REF, A only) and MTU-stretch-hold conditions (PL0, PL50, and PL95). Each colored dot represents a single participant. The thick and thin black horizontal and vertical bars indicate the group mean and standard deviation, respectively. ^#^ *p* < 0.01.

**Figure 5.**
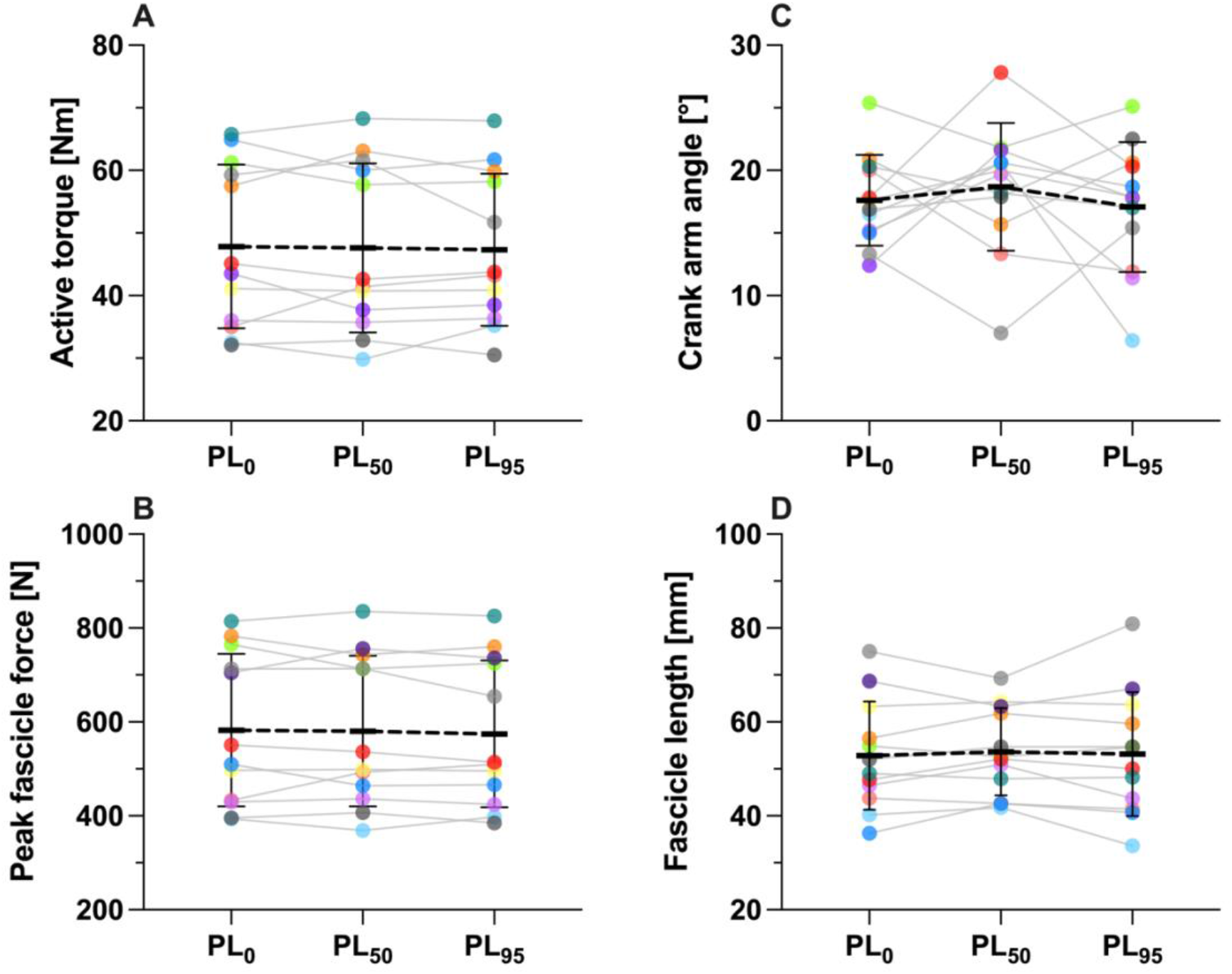
Individual (*n* = 12) and mean values of active torque (A), peak fascicle force (B), crank-arm angle (C) and fascicle length (D) at peak fascicle force during the crank-arm rotation across the three MTU-stretch-hold conditions (PL0, PL50, and PL95). Each colorful dot represents a single participant. The thick and thin black horizontal and vertical bars indicate the group mean and standard deviation, respectively.

### Peak torque during stretch

Peak torque was similar among the MTU-stretch-hold contractions (48.5 ± 13.2 Nm; *p* = 0.682; Figure 6A). Muscle activity (i.e., EMG envelope) was similar among the MTU-stretch-hold contractions at peak torque (0.579 ± 0.209 V, *p* = 0.463), but the crank-arm angle, fascicle length and fascicle velocity at the instance of peak torque were significantly different among the MTU-stretch-hold contractions (*p* ≤ 0.002; Figure 6 and Table 2). Fascicle force at the instant of peak torque was not significantly different among the MTU-stretch-hold contractions (*p* = 0.086).

**Figure 6.**
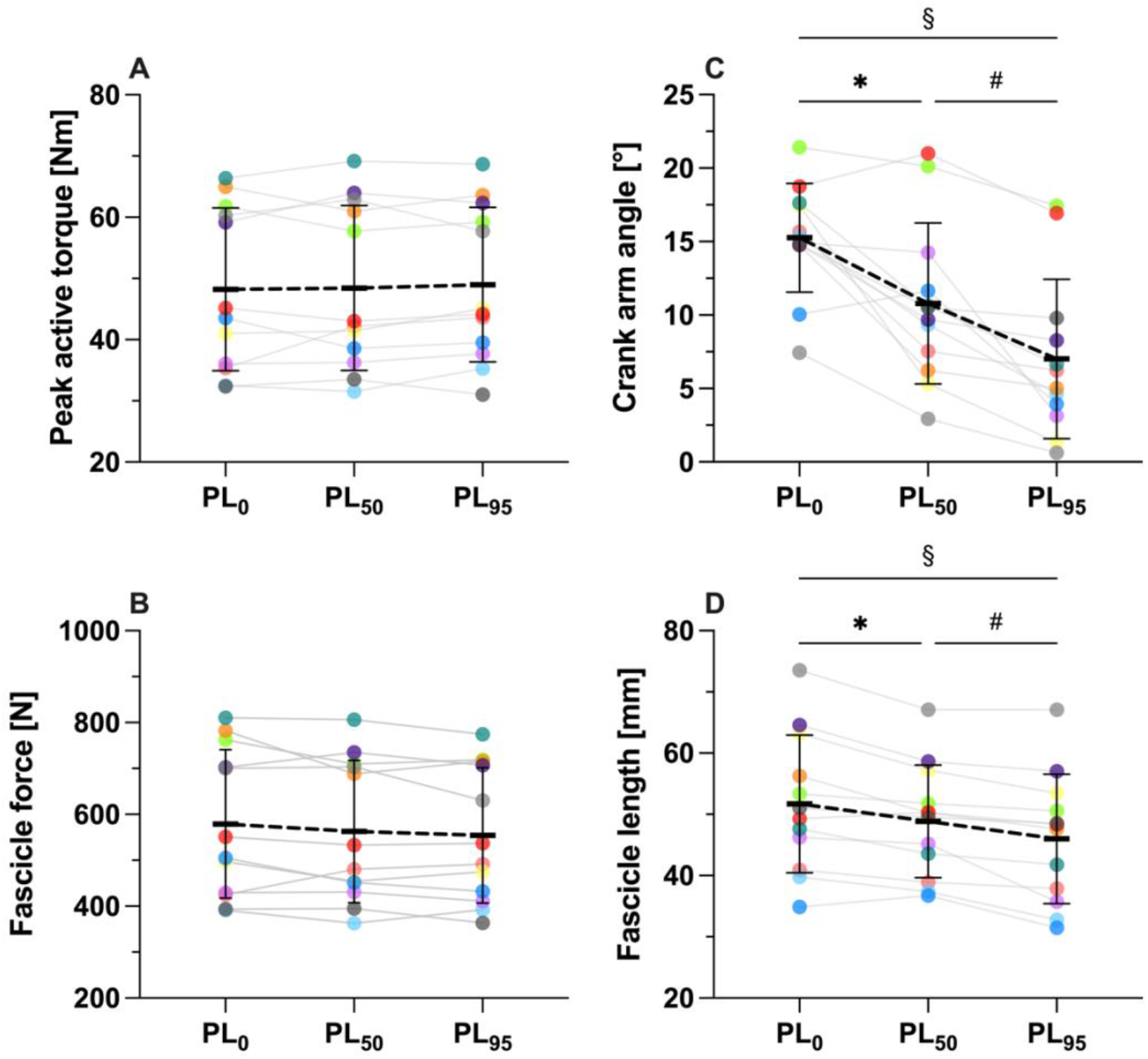
Individual (*n* = 12) and mean values of peak torque (A), fascicle force (B), crank-arm angle (C), and fascicle length at peak torque (D) across the three MTU-stretch-hold conditions (PL0, PL50, and PL95). Each colorful dot represents a single participant. The thick and thin black horizontal and vertical bars indicate the group mean and standard deviation, respectively. * *p* < 0.05; ^#^ *p* < 0.01; ^§^ *p* < 0.001.

**Table 2.**
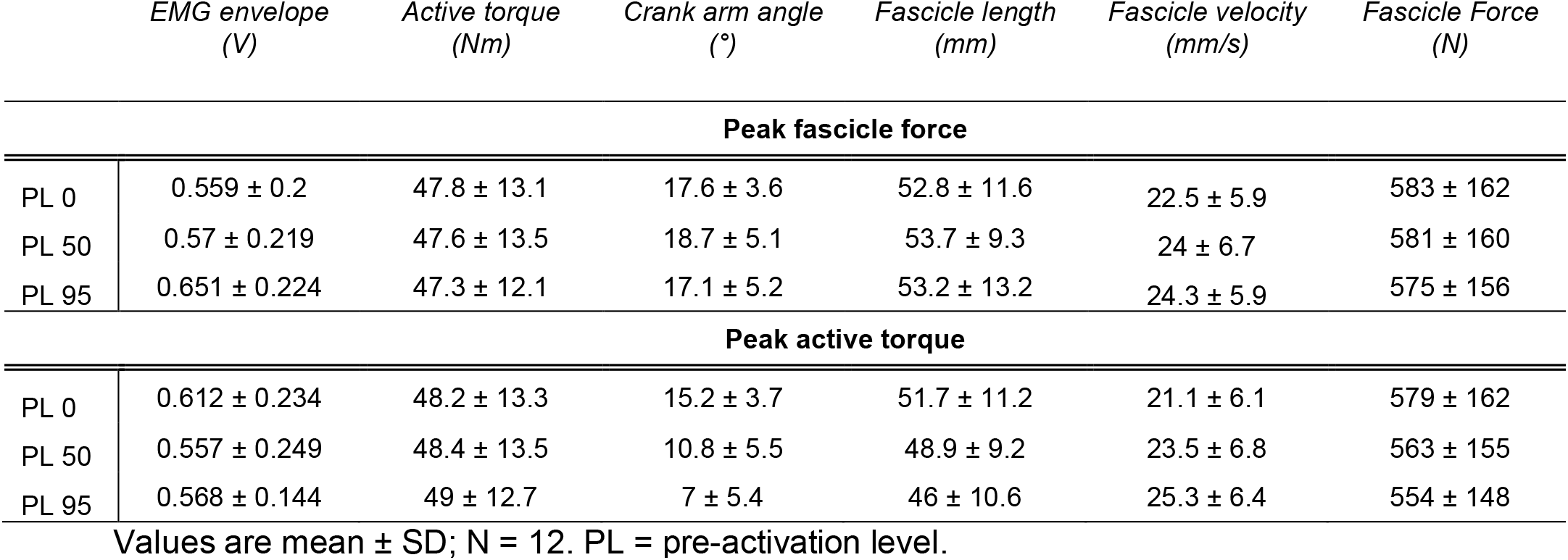
EMG envelope, active torque, crank arm angle, fascicle length, fascicle velocity and fascicle force values at peak fascicle force (top panel) and at peak active torque (bottom panel) across the three MTU-stretch-hold contractions.

### Residual force enhancement (rFE)

Steady-state torques were greater in all MTU-stretch-hold contractions compared with the REF condition (29.3 ± 7.2 Nm vs. 27.3 ± 6.1 Nm, *p* < 0.001), however the amount of rFE was similar among the MTU-stretch-hold contractions (7.0 ± 5.3% REF_TOR_, *p* = 0.559; Figure 4B). Steady-state fascicle lengths were similar among the contraction conditions (65.5 ± 12.1 mm, *p* = 0.304).

### Repeated measure correlations

No significant repeated-measures linear relationships were found between fascicle stretch amplitude to the peak fascicle force and fascicle force (r_rm_(23) = −0.41, 95% CI: −0.69 to −0.02, adjusted *p* = 0.125), and between fascicle stretch amplitude and rFE (r_rm_(23) = −0.03, 95% CI: −0.42 to 0.37, adjusted *p* = 2.649). A significant repeated-measures linear relationship was found between peak fascicle force during rotation and rFE (r_rm_ (23) = 0.62, 95% CI: 0.30 to 0.82, adjusted *p* = 0.003; Figure 7).

**Figure 7.**
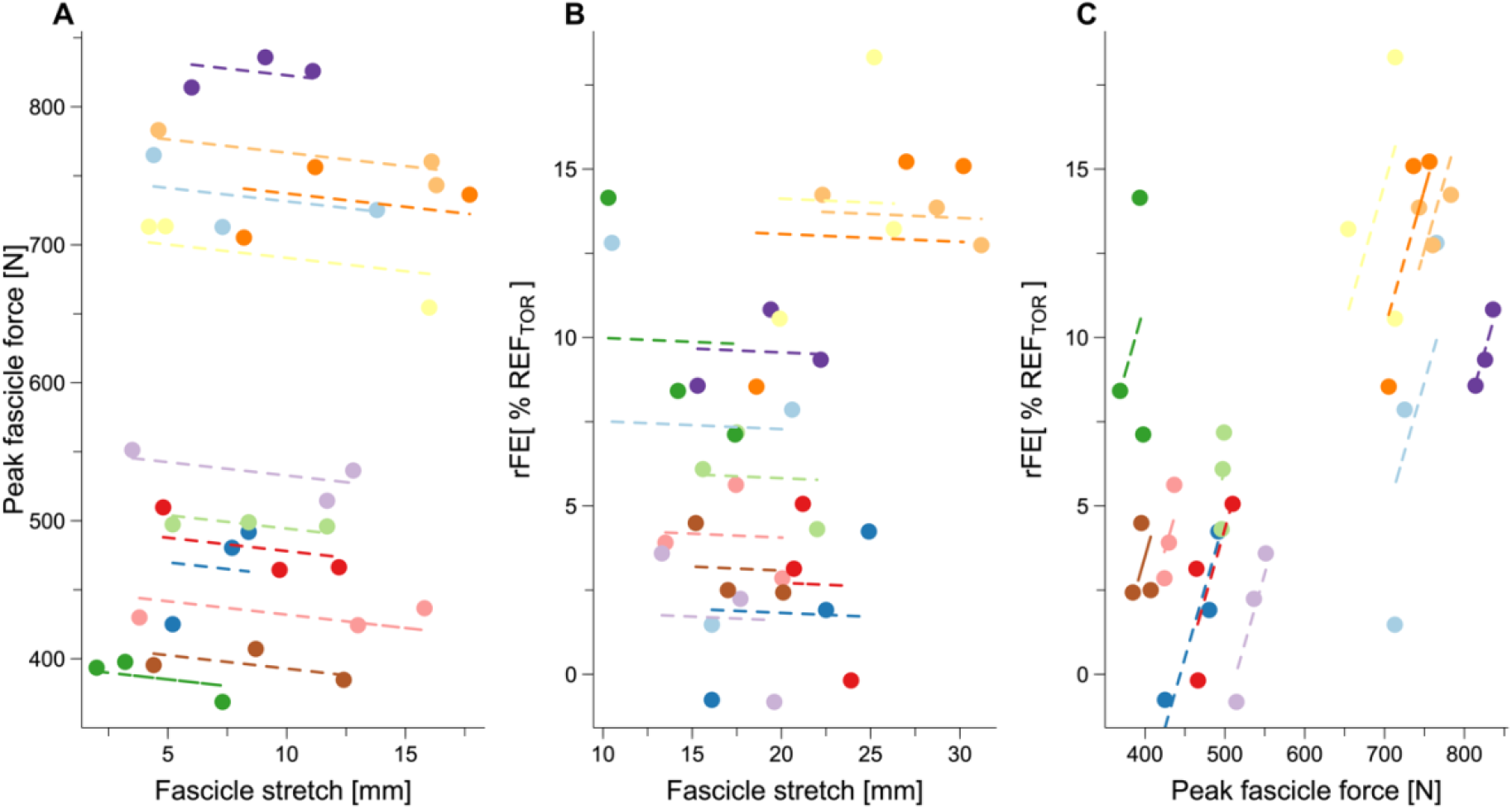
Repeated-measures linear relations between tibialis anterior muscle fascicle stretch amplitude until peak fascicle force and peak fascicle force (A: r_rm_(23) = −0.41, 95% CI: −0.69 to −0.02, *p* = 0.125, adjusted *p* value), fascicle stretch amplitude and rFE (B: r_rm_(23) = −0.03, 95% CI: −0.42 to 0.37, *p* = 2.649, adjusted *p* value), and peak fascicle force and rFE (C: r_rm_ (23) = 0.62, 95% CI: 0.30 to 0.82, *p* = 0.003). Each color represents a single participant (*N* = 12).

## Discussion

The purpose of this study was to quantify how changes MTU series elasticity from different pre-activation timings and levels (PLs) affects muscle fascicle kinematics and kinetics during apparent eccentric exercises. Despite similar joint kinematics, we found significant differences in fascicle kinematics during MTU-stretch-hold contractions, with higher PLs resulting in larger shortening and stretch amplitudes within the human TA (Figure 3, Table 1). This indicates that the PL substantially affects the fascicle strains experienced during apparent eccentric exercises, which could affect MTU adaptations to these exercises. The PL also affected fascicle work output, which was largely driven by length change differences rather than active force differences. These fascicle work differences could also affect morphological adaptations to eccentric exercise. Finally, despite different PLs and fascicle kinematics, the steady-state fascicle force following imposed MTU stretch was not significantly affected, which indicates the rFE was not dependent on fascicle stretch amplitude at the muscle length we investigated. Collectively, these findings indicate that changes in MTU elasticity from different PLs should be considered in eccentric exercise regimes because of its influence on fascicle kinematics and kinetics during MTU stretch.

We changed the PLs relative to the MTU stretch onset and observed that both fascicle shortening and stretch amplitudes increased with increasing PL (i.e., pre-activation), which supports our first hypothesis. Fascicles in PL0 shortened during, but not before the imposed MTU stretch. In PL50, fascicles shortened both before and during the MTU stretch. The greatest shortening amplitude occurred in PL95, where fascicles shortened before, but not during MTU stretch. Consequently, increasing the PL resulted in shorter fascicles at MTU-stretch onset. As the final MTU and fascicle lengths were similar among conditions, the fascicle stretch amplitudes during MTU stretch thus increased with PL. These results support previous findings on the human vastus lateralis and gastrocnemius medialis (31, 32) and can be explained by the relation between muscle activity level and tendon elasticity. Due to tendon elasticity (29, 30), fascicles shortened in PL50 and PL95 prior to MTU-stretch onset by stretching the tendon and other in-series elastic tissues. This results in a stiffer tendon (and other in-series elastic tissues) that can buffer less fascicle stretch during MTU stretch compared with when the same tendon is shorter and less stiff at MTU stretch onset. In PL95, the tendon presumably had the lowest capacity to buffer fascicle stretch because MTU elasticity was at its lowest among the conditions as the tendon was already stretched before MTU-stretch onset. In PL50, the tendon and in-series elastic tissues could buffer more fascicle stretch compared with PL95 because MTU elasticity was higher at MTU-stretch onset. In PL0, the muscle fascicles shortened the most during MTU stretch and the tendon and in-series elastic tissues were most effective at buffering fascicle stretch during MTU stretch (58). This is likely to be because MTU elasticity was at its highest among the conditions at MTU-stretch onset, and MTU elasticity took time to decrease as muscle activity and force rose during MTU stretch. Consequently, fascicle length changes depend on MTU series elasticity and muscle activity level as we showed, as well as MTU stretch velocity, similar to muscle-tendon interactions during pre-activated shortening contractions (59).

The increased PL along with the observed differences in fascicle kinematics led to both greater joint work and muscle fascicle work, which partly supports our second hypothesis. Joint work indeed increased by increasing PL, which was because of higher net joint torques at MTU stretch onset. Contrary to our hypothesis, positive fascicle work was not significantly different between PL0 and PL50, while it was significantly higher in PL95. The positive fascicle work differences were likely driven by differences in MTU elasticity at MTU stretch onset and the MTU lengths where maximal muscle activation occurred. In PL0 and PL50, maximal muscle activity was not attained until during MTU stretch and subsequently fascicle and MTU lengths were longer at maximal muscle activity compared with PL95. Due to a rising activity level during MTU stretch in PL0 and PL50, the fascicles were not able to shorten as much against in-series elastic tissues that were being stretched and becoming stiffer. However, in PL95, more fascicle shortening occurred at a fixed and shorter MTU length against higher MTU elasticity (43). Therefore, fascicle length changes during apparent eccentric exercises also depend on the MTU length range and the corresponding muscle activity changes.

In agreement with our second hypothesis, negative fascicle work increased linearly with increasing PL, which was largely because the fascicle stretch amplitudes increased with increasing PL. Notably, there was approximately a 20% difference (SPD) in negative fascicle work between PL0 and PL95. This difference in fascicle work is more than double the difference in joint work (9% SPD, Table 1) between the same two conditions. These findings suggest that eccentric exercise protocols that rely solely on external joint work as a workload index may substantially underestimate the energy absorbed by the muscle, as well as differences in muscle energy absorption among similar eccentric exercises. Differences in muscle energy absorption could subsequently affect muscle fascicle adaptations. However, it remains unclear whether the key mechanical factor driving specific fascicle adaptations to eccentric exercise is the work performed by the fascicle or rather its absolute length change, the fascicle lengths experienced during the length change, or the mean or peak fascicle force produced during MTU stretch.

We observed that similar peak fascicle forces occurred at similar fascicle lengths and joint angles irrespective of the PL and fascicle stretch amplitudes, supporting our third hypothesis. As we also observed similar muscle activity levels, but different fascicle stretch amplitudes until the peak fascicle force, it seems that the peak fascicle force was determined by the muscle’s active force-length relation because the lengths at peak fascicle force coincided with the plateau region of this relation (43, 60). This interpretation is supported by our observation that fascicle forces did not increase further, but decreased when the stretches continued onto the descending limb of the TA’s active force-length relation. Unlike peak fascicle force, peak ankle joint torque occurred at varying joint angles, fascicle lengths, and fascicle velocities (see Table 2). This raises intriguing questions regarding Herring’s notion (61) that muscles tend to adapt in the “active position”. As peak fascicle forces were independent of PL and thus similar across the eccentric exercise conditions, other factors (e.g., fascicle work or stretch magnitudes) might better explain the variability in muscle fascicle adaptations to eccentric exercise.

Although we observed different fascicles kinematics and kinetics, rFE was relatively constant across the three MTU-stretch-hold conditions, which supports our fourth hypothesis. Our rFE values (6.5-7.6%, Figure 4B) are comparable with those previously reported from the human dorsiflexors (62–64). Notably, fascicle stretch amplitude did not affect rFE (Figure 7A), which contradicts previous findings that showed higher rFE with larger stretch amplitudes (32, 35, 65). However, this previous study found differences in peak torques during MTU stretch (32). Our findings align with those of from Hisey et al. and Bakenecker et al. (40, 66), who showed that rFE depends on the final muscle length rather than the fascicle stretch amplitude. Additionally, our findings showed that rFE strongly depends on the peak fascicle force during stretch (Figure 7C) as similarly reported by Bullimore et al. (33) for the in-situ cat soleus and by Paternoster et al. (36, 37) for peak torque during in vivo human studies. However, using peak torque instead of peak fascicle force makes it challenging to draw clear conclusions related to rFE because similar peak torques occurred at different joint angles, MTU lengths, and fascicle lengths in our study (Figure 6, Table 2).

Our rFE findings further suggest that a passive element arranged in parallel with the active cross bridges may be engaged at a specific fascicle length during MTU stretch, which could be the length where passive muscle force naturally increases, or the optimal fascicle length. Due to progressive fascicle stretch, this passive element could have progressively increased its force contribution to rFE once it was engaged. We speculate that this passive element, possibly titin (67), was not engaged upon muscle activation at the initially short lengths in our MTU-stretch-hold conditions, but that the passive element became engaged during active muscle stretch around the same muscle length, which resulted in it being stretched similarly across conditions. Subsequently, its passive force contribution remained similar following stretch across our MTU-stretch-hold conditions, which was reflected by similar rFE. We further speculate that terminating the stretch at a longer final fascicle length might lead to greater rFE because of greater stretch of passive elements. However, stretching from a longer initial fascicle length beyond the length of peak fascicle force would potentially induce less rFE because of reduced passive force contributions from passive elements.

Although we implemented MTU-stretch-hold and fixed-end reference contractions, fascicles clearly shortened in each contraction upon muscle activation due to MTU series elasticity. Consequently, residual force depression (rFD), which is induced by active fascicle shortening (68), might have affected the force output during the contractions. Previous studies that investigated shortening-stretch contractions of cat soleus muscle, which were similar to the fascicle behavior we observed, showed that rFE following such contractions was attenuated or even abolished (69, 70), which has been attributed to the muscle work performed during shortening (71). Yet, our findings showed relatively constant peak fascicle forces during stretch and rFE following stretch across the different conditions, regardless of fascicle shortening amplitude and the positive fascicle work. This is in line with a recent study that showed that rFD of the in vivo TA muscle is not related to positive fascicle work (72). Consequently, we think that the fascicle stretch amplitudes in our contractions were large enough that the potential rFD induced at the start of the contractions was abolished. However, our magnitudes of rFE might be overestimated because of rFD contamination in the fixed-end reference contractions (72, 73).

### Implications for eccentric exercise design

Eccentric exercise can be manipulated by changing the activation relative to MTU stretch onset (i.e., pre-activation), or by changing the amount of muscle activation during MTU stretch, which affects muscle fascicle mechanics. The insight provided by this study is important for designing targeted training interventions to enhance muscle performance, improve rehabilitation outcomes, and prevent injury. For example, we showed that increasing the PL led to more joint and muscle fascicle work, and consequently a high PL might be preferred in eccentric exercise protocols that aim to limit the number of repetitions while attempting to maximize muscle adaptation. Additionally, a higher PL increased fascicle shortening amplitudes, resulting in increased tendon strain throughout MTU stretch. Tendon strain is thought to drive the tendon’s anabolic response and its subsequent adaptations (74), which suggests that an increased PL might be more beneficial for treating or preventing tendinopathy. Finally, our previous (40) and current rFE findings suggest that peak fascicle force during MTU stretch and the final muscle length the muscle is stretched to during eccentric exercise might be crucial for inducing specific adaptations that titin regulates such as longitudinal muscle hypertrophy (41).

## Limitations

A few pitfalls need to be considered regarding the TA fascicle force estimates. First, we used literature-based TA moment arm values and extrapolated them using a second-order polynomial fit (54) over −15° DF to +30° PF (Figure S2). However, the TA moment arms we used, which were determined during maximal voluntary dorsiflexion contractions (add reference), did not correct for potential talus rotations, which Miller et al. (75) found to affect TA moment arm values at rest. The absolute values we reported may be affected by TA’s tendinous insertion location, which we assumed was constant between participants, but this is likely to not be the case (71,72) and could affect TA’s function and force-generating capacity. We assumed TA’s force contributions were equivalent among the conditions based on its similar activity levels, but it is possible that pre-activation affected its relative force contributions during the contractions. Further, we are aware that anatomical cross-sectional area (ACSA) is not a good predictor of muscle force generation capacity (76), yet Nagayoshi et al., (77) showed that the maximal ACSA seems to be an adequate predictor of the torque generation capacity of the dorsiflexors. This can be explained by the small fascicle angle variations we found. Indeed, the difference in fascicle angles we observed during the contractions between participants and conditions was 12.4° on average, and the 95% CI was from 11.8° to 13.0°. The maximum fascicle angle we found was 22.8° (Figure S4). Hence, a minimum of ~92% of the force generated is theoretically transmitted longitudinally (the cosine of 22.8°). Given the small fascicle angles observed, ACSA appears to be a reasonable estimator of force for the TA muscle fascicles. Based on previous findings (78, 79), negligible co-contraction was assumed over the entire ROM. Further, crank arm angle was assumed to reflect joint angle as previous studies defined ankle angle between the tibia and sole of the foot (56). However, as we used a repeated measure study design our conclusions should not be largely affected by these limitations.

## Conclusions

Our study sheds light on fascicle mechanics variations during apparent eccentric exercise. We found that eccentric exercise always resulted in fascicle shortening followed by fascicle stretch rather than isolated eccentric contractions. Such fascicle behavior is caused by the MTU’s series elasticity and changes in this elasticity because of alterations in the timing and level of muscle activation during eccentric exercise. As the resulting fascicle mechanics differed across the eccentric exercise conditions, eccentric exercise that appears to be similar at the joint and MTU levels might trigger different muscle fascicle adaptations, which might explain the large variability in adaptations reported following apparent eccentric exercise. Accordingly, it is crucial to understand the underlying fascicle mechanics, which should be considered and standardized when eccentric exercise protocols are designed and evaluated. Future studies should investigate which fascicle mechanics are key in driving specific adaptations to apparent eccentric exercise.

## Data availability

The scripts used for the data collection and analysis, along with the data and extra medias, are freely available at the following GitHub repository: https://github.com/PaulT95/Tecchio_et_al_2023.

## Acknowledgements

We would like to thank all of the participants for donating their time.

## Author Contributions

PT, BJR and DH conceived and designed the research. PT performed the experiment, analyzed data, prepared the figures, drafted the manuscript. PT, BJR and DH interpreted results, edited and revised the manuscript. All authors approved the final version of the manuscript.

## Supplemental Material

**Figure S1.**
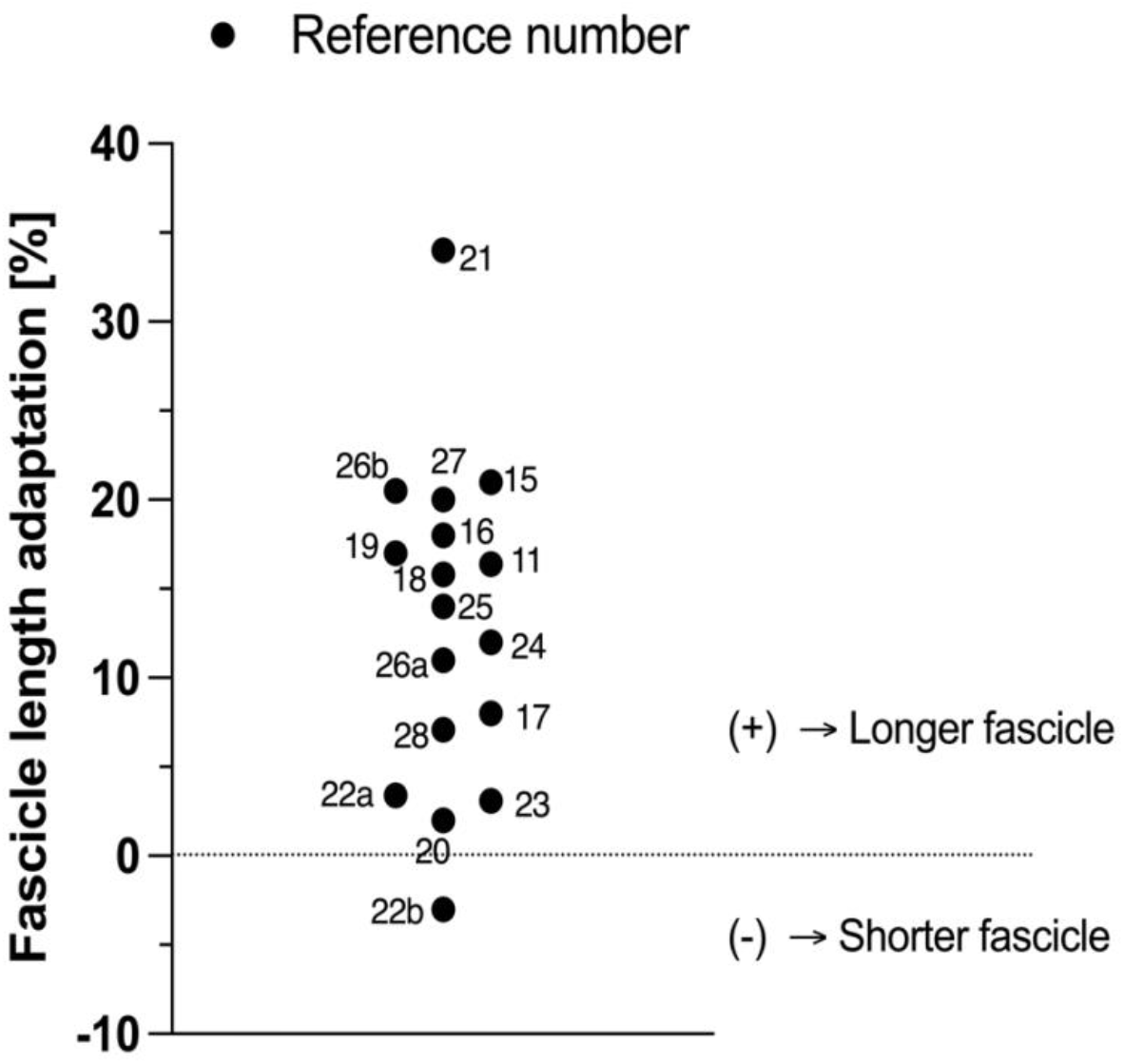
Fascicle length adaptations following eccentric exercise of lower limb in vivo human muscles reported in the literature. The numbers next to the black dots represent the reference number of the study.

**Figure S2.**
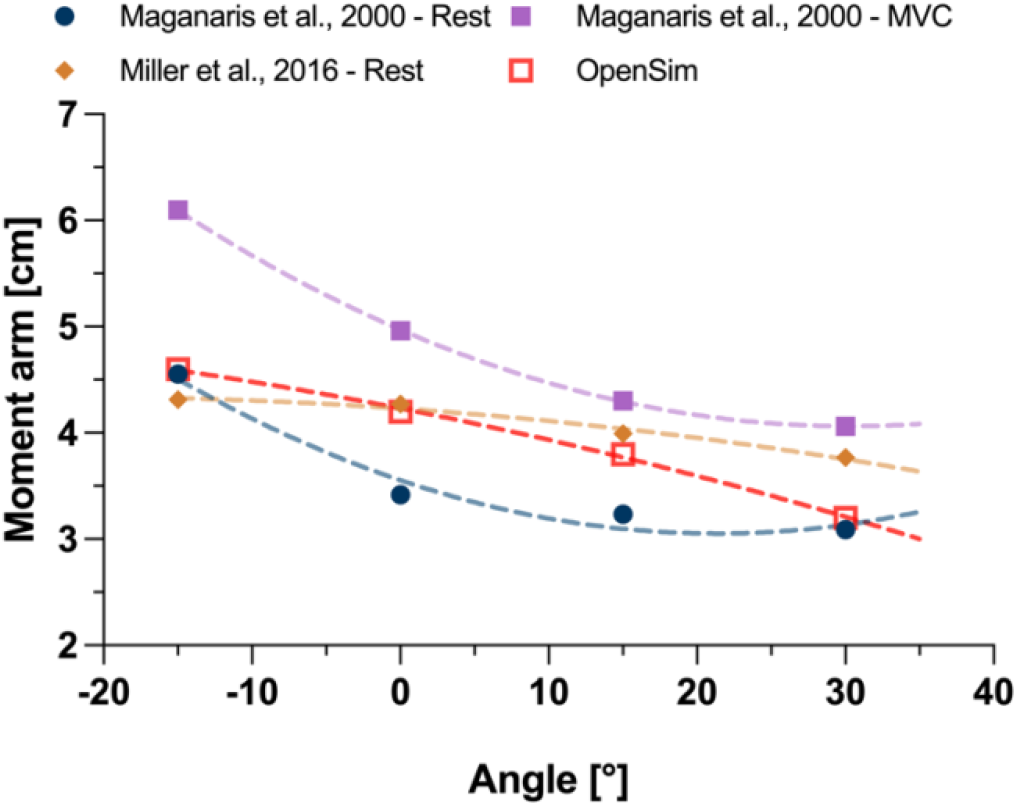
Tibialis anterior moment arm values measured from three studies and estimated in OpenSim at four ankle joint angles relevant for this study (−15° dorsiflexion, 0°, 15° and 30° plantarflexion) at rest (2 studies) and during maximal voluntary contraction (1 study). The dashed lines represent second-order polynomial fits that were extrapolated to +35° plantarflexion.

**Figure S3.**
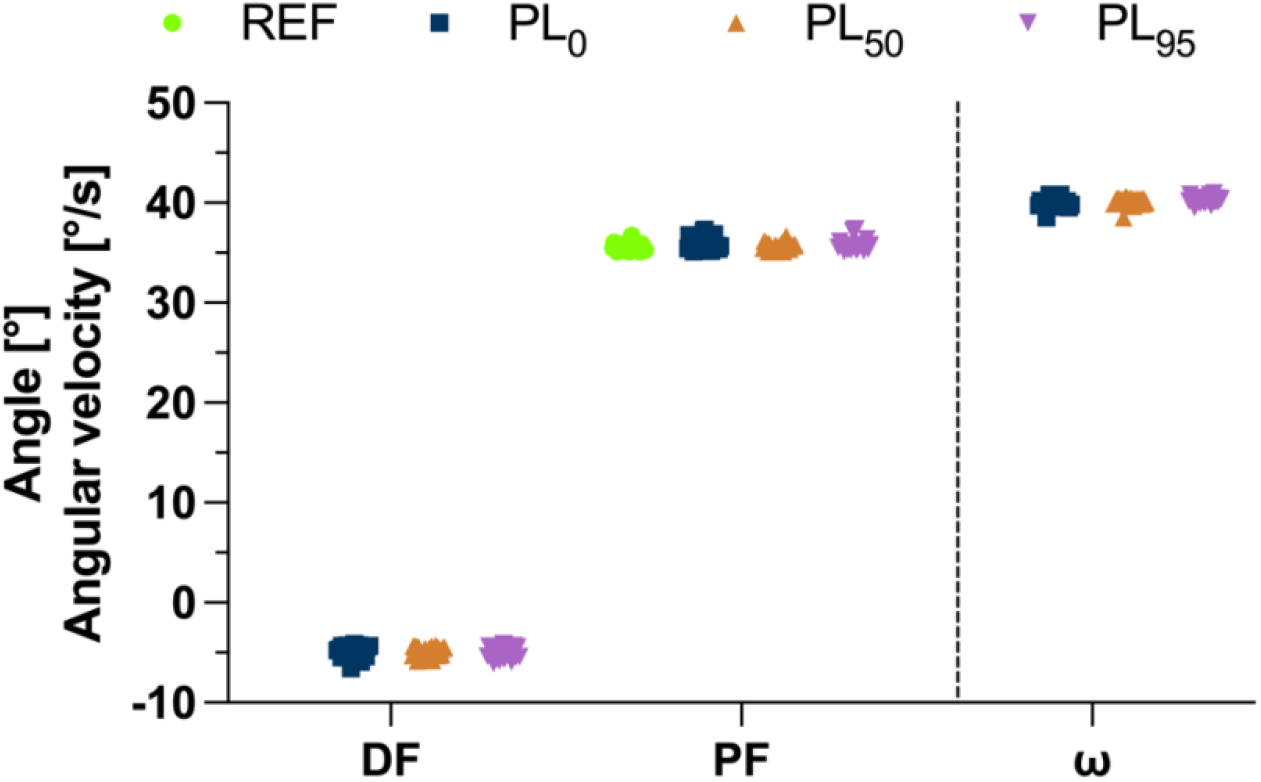
Individual (*n* = 12) crank arm angle values at the dorsiflexed position (−5°, DF) and plantarflexed position (+35°, PF) across the different contraction conditions, along with the mean angular velocity (ω) during the three MTU-stretch-hold conditions.

**Figure S4.**
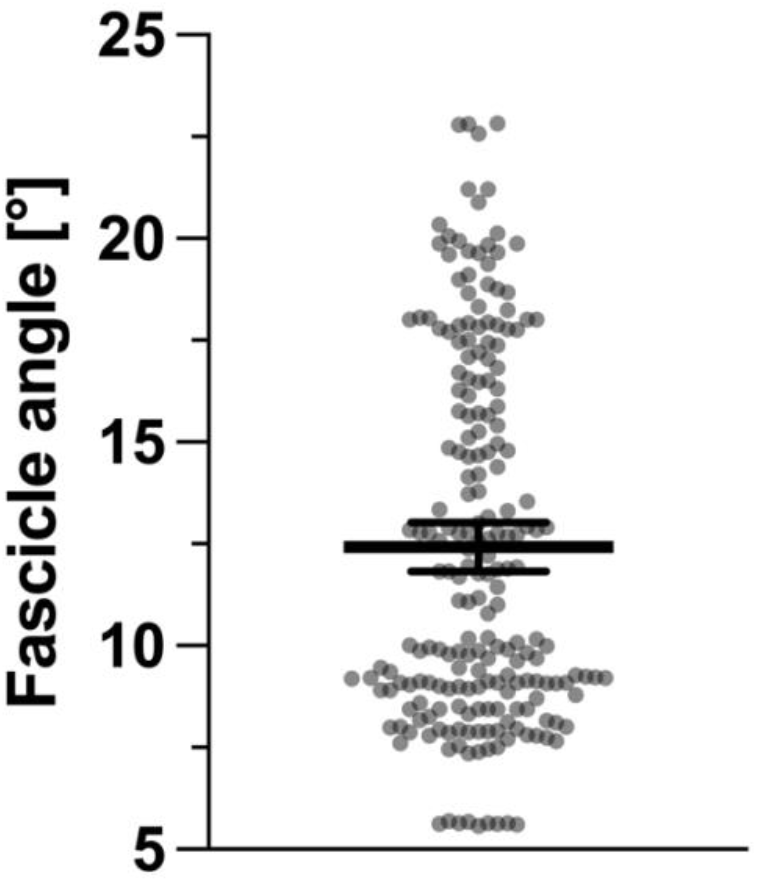
Tibialis anterior superficial compartment fascicle angle variations from rest to maximal voluntary contraction over a 40° range of motion across 12 participants. The gray dots represent single values of the participants’ minimum and maximum values during the conditions, and the thick and thin black horizontal lines represent the mean and the 95% confidence intervals, respectively. Mean = 12.4°, SD = 4.3, 95% CI = 11.8° - 13.0°.

